# MaCADEMIA: a software toolbox for single-molecule fluorescence image quantification

**DOI:** 10.64898/2026.10.30.763064

**Authors:** Dominic Alderson, Zoe Catchpole, Katrina Crompton, Katherine Sanders, Sophia Johnson, Kirsty Smith, John Packer, Kate S. Harris, Josana Rodriguez, Mark C. Leake, Adam J. M Wollman

## Abstract

Single-molecule localisation microscopy (SMLM) is a powerful tool in molecular and cellular biology and part of the broad umbrella of super-resolution microscopy. A distinct feature of SMLM, is its capacity for deep quantification, beyond just spatial information, to copy number, stoichiometry, molecular interaction and dynamics in living cells. The hardware required is increasingly straightforward to implement. However, complexity remains in the image analysis where much remains fragmented across specialised software tools. Here, we present MaCADEMIA (Matlab Computational Analysis of Diffusion, Emitter Molecular Intensity and Abundance), a simple yet powerful toolbox of MATLAB functions and associated documentation to enable straightforward extraction of quantitative single-molecule information. MaCADEMIA performs ensemble particle detection achieving localisation accuracies down to 0.5 pixels Root mean squared error (RMSE) across wide signal-to-noise (SNR) ranges on standardised particle tracking benchmarks, consistently matching or exceeding the performance of 14 leading methods. Our integrated workflow enables molecular counting and stoichiometry determination through stepwise photobleaching analysis, mobility measurement via mean squared displacement, colocalisaton analysis and absolute protein copy number quantification using three-dimensional cellular models.

## Introduction

Single-molecule localisation microscopy (SMLM) encompasses a broad range of imaging techniques. Each of these has the sensitivity to detect individual fluorescently tagged molecules and overcome the optical diffraction limit to precisely determine the location of each molecule. SMLM comprises two broad, complementary approaches: super-resolution reconstruction and single particle tracking (SPT) – Figure 1. In super-resolution reconstruction, the localised positions are used in a ‘pointillist’ approach to reconstruct highly detailed super-resolved images, e.g. Photo-Activated Light Microscopy (PALM) and Stochastic Optical Reconstruction Microscopy (STORM)^1,2^. In SPT, single emitters or complexes of emitters are super-resolved and tracked over time. By careful analysis, a wealth of information can be extracted from SPT. The position of single-particles over time can reveal the behaviour of biological molecules in cells, such as the treadmilling of FtsZ involved in the division of bacterial cells^3^. Time-resolved tracked locations can be used to characterise diffusive motion, for instance revealing whether transcription factors are bound or unbound in the nucleus.^4,5^ SPT can also be used for molecular counting, where the number of molecules within a complex can be determined by quantifying the intensity of fluorescently-tagged particles as they photobleach.^6,7^ Molecular counting has been used to determine the architecture of the DNA replication machinery in bacterial^8^, and the carbon fixing, carboxysome in photosynthetic bacteria.^9^ SPT can be simultaneously performed in multiple colour channels, also quantifying molecular and protein-protein interactions.^10^ Together, SMLM+SPT has provided key insights into the molecular architecture of complexes *in vitro*^11^ and inside live prokaryotic^10,12,13^ and eukaryotic (including human) cells.^14–16^

**Figure 1:**
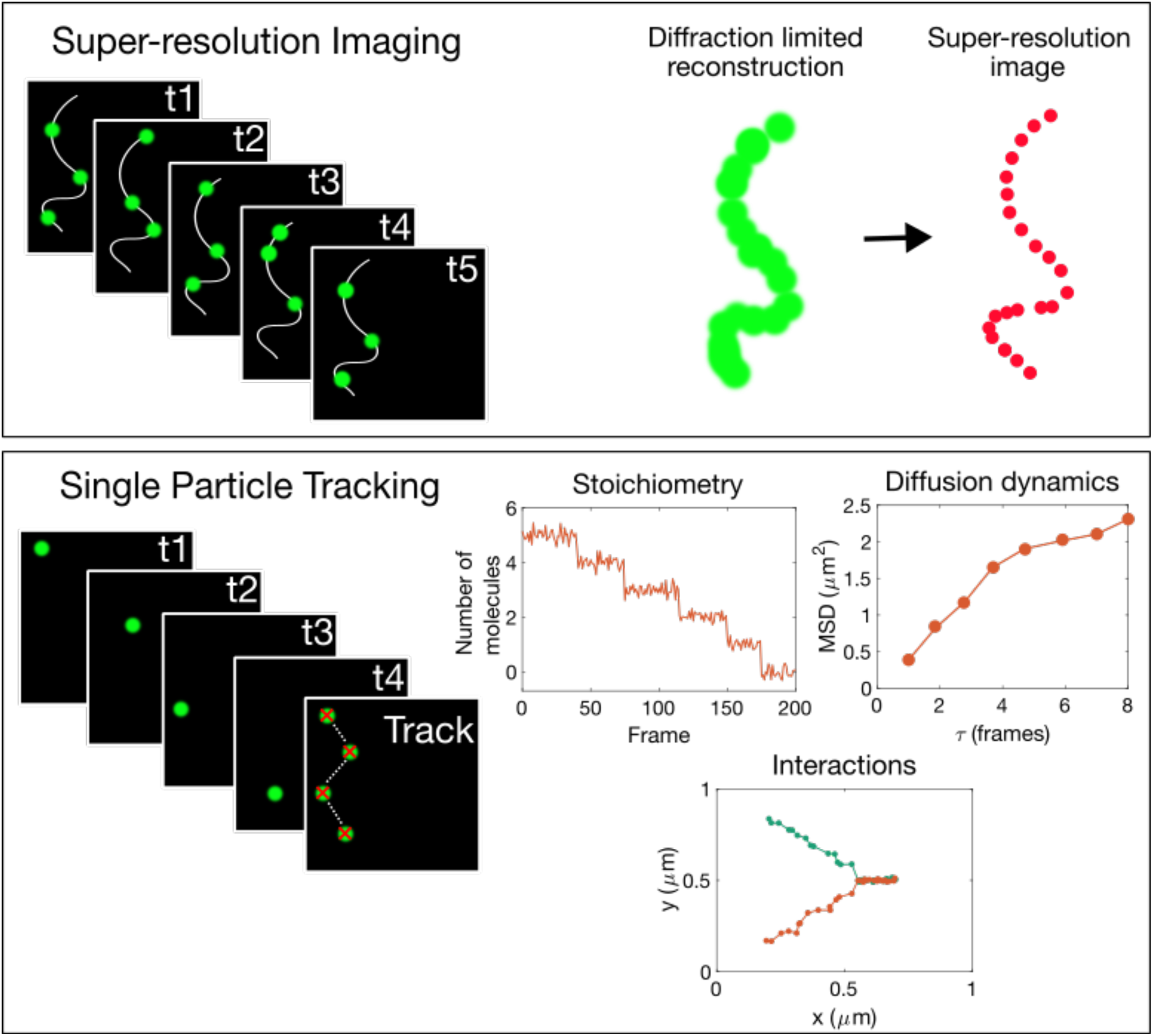
Complementary single-molecule ffuorescence microscopy modalities. (Top) Super-resolution imaging accumulates localisations across many frames (t1 to t5) to reconstruct structures below the diffraction limit, replacing the diffraction-limited image (green) with a pointillist map of emitter positions (red). (Bottom) Single particle tracking instead links localisations of the same molecule across frames to recover trajectories, yielding multiple per-molecule observables: stoichiometry from stepwise photobleaching of intensity I in units of number of single molecule equivalents (left top), diffusion dynamics from the MSD versus lag time τ (right top), and molecular interactions from co-trajectories of two spectrally distinct labels (bottom).

From a pattern recognition perspective, robust single molecule analysis requires the solving of multiple interconnected problems under challenging conditions. Particle detection can be considered a binary classification problem where signal-to-noise ratios (SNR) often fall below the 2:1 range requiring methods that preserve sensitivity whilst controlling false positive rates^17,18^. Individual detection approaches exhibit complementary strengths. Single-molecule detection at low SNR presents fundamental challenges in balancing sensitivity and specificity, with detection performance varying notably across different imaging conditions and molecular dynamics^19^. Trajectory linking requires temporal pattern matching across frames with stochastic intensity variations and sporadic observations. Segmentation demands boundary detection in heterogeneous images where deep learning approaches like Cellpose^20^ require extensive training data. The fundamental challenge lies in maintaining robust performance across these diverse pattern recognition tasks within a unified, interpretable framework.

SMLM relies on computationally fitting the point spread function (PSF) generated by individual molecular complexes. To aid researchers in analysing this data, numerous algorithms and software packages have been developed, either as standalone suites or as plugins for programs like ImageJ, including the popular TrackMate plugin.^21^ However, the existing software landscape is highly specialised. Many popular packages, such as the versatile ThunderSTORM ^22^, are designed for the high-precision reconstruction of static, super-resolved images. Other widely used algorithms like RainSTORM ^23^, QuickPALM ^24^, and DAOSTORM ^25^ also offer high spatial resolution but lack a focus on temporal information. DAOSTORM, for example, excels at handling images with a high density of fluorophores, while QuickPALM includes utilities for 3D reconstruction and drift correction valuable for complex workflows.^26^Some algorithms incorporate pre-filtering steps such as Gaussian filtering ^27–30^, Laplacian of Gaussian ^28,29,31^, wavelet products ^32,33^, or deconvolution ^34^.More innovative routines utilise machine learning, like 3B (Bayesian analysis of bleaching and blinking)^35^ or similar AI/ML approaches such as Kalman filtering or CNNs^36–38^. Although these methods promise efficient analysis of large datasets, they can function as “black-boxes,” requiring careful interpretation of results and thoughtful selection of models and priors.

Despite this ecosystem of specialised tools, the fragmented software landscape creates a significant hurdle for researchers. Scientists focused on quantitative live-cell analysis are often forced to create cumbersome workflows, using one tool for localisation, another for tracking, and custom scripts for essential pre-processing tasks like cell segmentation. This not only presents a steep technical barrier but also introduces multiple potential points of error. TrackMate can unify tracking and segmentation^21^ but it lacks the capability for deep quantification in SMLM data, including stoichiometry and copy number determination. There is a need for a single, integrated software package that combines robust live-cell tracking with user-friendly stoichiometry determination and other essential pre-processing tasks like cell segmentation. This highlights a need for transparent, non-black-box software solutions. An algorithmic approach ensures that the entire workflow is tuneable, allowing researchers to optimise parameters, understand their effects on the results, and validate their analytical pipeline against ground truth data; a step that is often opaque in machine learning/AI based solutions.

MaCADEMIA is a comprehensive, MATLAB-based software package designed to make advanced single-molecule analysis accessible to a broader range of researchers. It provides a complete suite of tools to quantify key physical information from microscopy data, including cell segmentation, single-particle tracking, protein stoichiometry, diffusion coefficients, and cellular protein copy number. Alongside its analysis capabilities, MaCADEMIA features a powerful simulation module for generating the ground-truth data required for algorithm benchmarking.

While Python is a growing force in scientific computing, the choice of MATLAB for MaCADEMIA was an informed one, leveraging its extensive and professionally maintained built-in image analysis functions. This rich, integrated environment allows for the rapid development of reliable and complex analysis tools without needing to build foundational components from scratch. To ensure the underlying methods are accessible to the entire research community, we have also supported development of PySTACHIO ^39^, a complementary open-source Python implementation of the tracking and stoichiometry functions. Additionally, MaCADEMIA supports integration with OMERO enabling compatibility with modern image data management infrastructure^40^. Ultimately, MATLAB provides the ideal platform for building MaCADEMIA as a single, cohesive, and powerful software package.

## Results

### MaCADEMIA Single particle tracking

To identify and quantify diffraction-limited fluorescent foci, hereafter referred to as “spots”, MaCADEMIA employs configurable, automated detection methods (see Methods for full details). By default, the software applies a Laplacian of Gaussian (LoG) filter to each frame to enhance spot-like features of a specified radius, followed by an adaptive thresholding step based on image statistics. Alternatively, a bespoke morphological detection algorithm can be used for extremely low SNR environments. Following initial detection and noise-removal steps, each potential spot is localised to sub-pixel precision using an iterative Gaussian-weighted centroid algorithm.^32,41^ Finally, the intensity for each spot is determined by summing the pixel values within its inner radius and subtracting a locally calculated background value.

To quantitatively benchmark our algorithm, we adopted the standardised methodology established by the open Particle Tracking Challenge, which objectively compared multiple tracking codes on simulated datasets^42^. Our software, MaCADEMIA, was evaluated using the benchmark datasets for simulated particles undergoing Brownian and confined diffusion across a range of signal-to-noise ratios (SNR) and particle densities (Supplementary figure 1). MaCADEMIA’s output was compared against the provided ground truth, with a true positive defined as a detection within a 3-pixel radius of a true particle. Performance was quantified by two key metrics: Jaccard Index (J) and Root Mean Square Error (RMSE). MaCADEMIA demonstrated robust performance, with both accuracy and detection success improving with higher SNR and lower particle density as expected (Figure 2). When compared to the 14 methods tested in the open particle tracking challenge benchmark^42^, our algorithm proved highly competitive. Localisation RMSE fell to 0.3-0.5 px at SNR 7, and across every tested density it tracked the lower, more accurate edge of the published range, matching the best-performing algorithms and falling at or below their lower bound at high SNR (Figure 2D to F, Supplementary figure 2). This validates the effectiveness of our centroid-based sub-pixel refinement approach.

**Figure 2:**
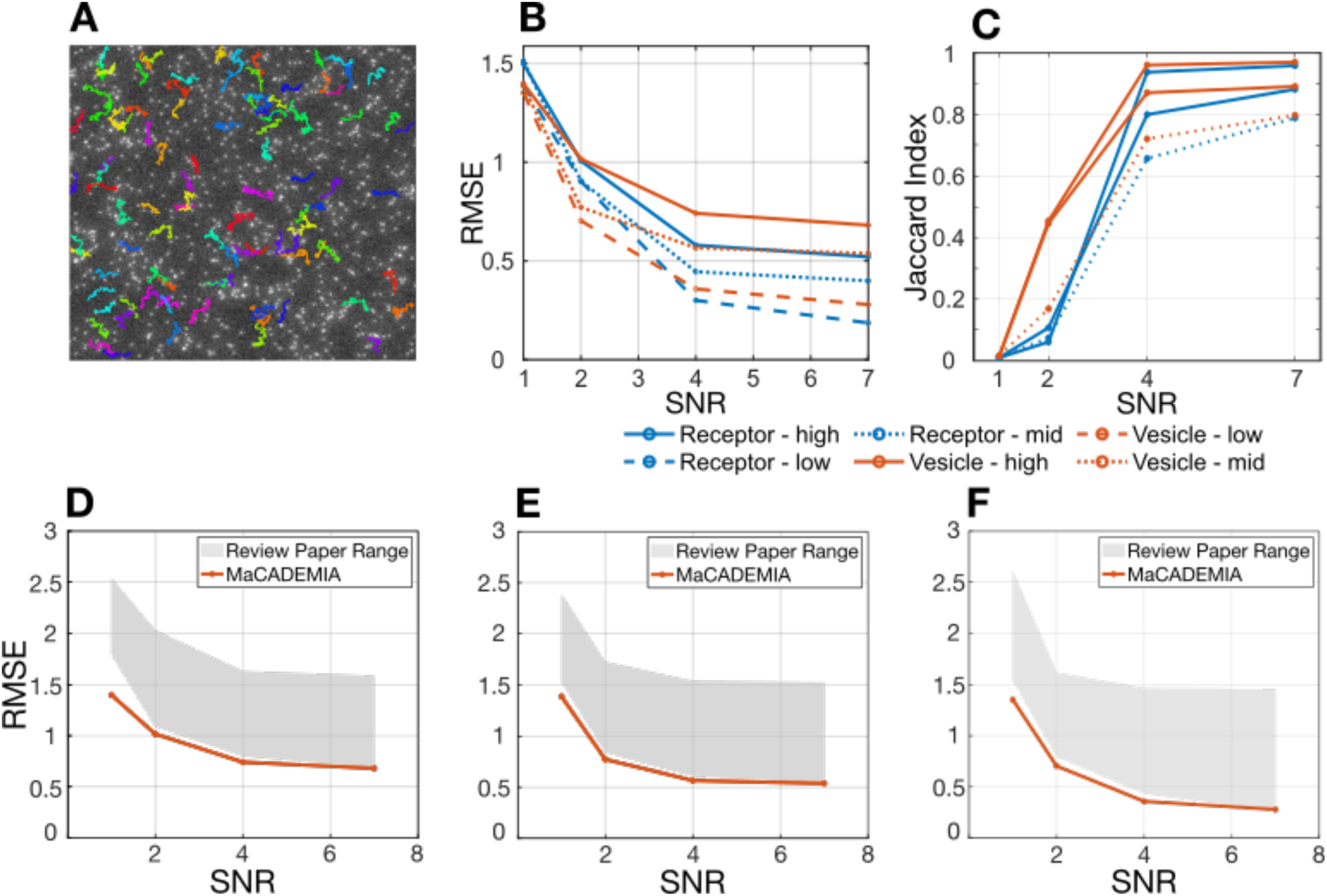
Tracking performance across varying signal-to-noise ratios and particle densities. Evaluation of the MaCADEMIA algorithm using benchmark simulated datasets for receptors (confined motion) and vesicles (Brownian motion). **(A)** Example single frame from vesicle simulation (SNR=4, high density) with ground truth trajectories overlaid in colour, showing the top 100 highest-displacement particles. **(B)** Localisation accuracy measured by Root Mean Square Error (RMSE) in pixels across SNR conditions. RMSE values range from approximately 1.5 pixels at SNR=1 down to 0.5 pixels at SNR=7, with vesicle datasets showing slightly higher error than receptor datasets due to increased particle mobility. **(C)** Detection success quantified by the Jaccard Index, where a score of 1.0 indicates perfect detection. Performance ranges from 0.7 to 0.85 depending on particle motion type and density, demonstrating robust detection that improves with higher SNR and lower particle density. **(D-F)** Localisation error (RMSE) versus SNR for vesicle datasets at high (D), medium (E), and low (F) particle densities. MaCADEMIA (orange line) achieves RMSE of 0.3–1.5 pixels across the SNR range 1–7, consistently matching or exceeding the performance range of peer-reviewed algorithms (grey shaded region), validating the centroid-based sub-pixel refinement approach. Note that for RMSE, lower values indicate superior accuracy.

MaCADEMIA processed the high-density vesicle (SNR 7) benchmark sequence in 18.2 s, equivalent to 5.5 frames per second. Under identical conditions, TrackMate^21^ using the LoG detector with LAP linking completed the same task in 3.1 s (1.4 s detection, 1.7 s tracking; approximately 33 fps), a six-fold throughput advantage. This difference reflects the sub-pixel localisation step rather than detection. Both pipelines use a Laplacian-of-Gaussian filter for initial spot identification; however, MaCADEMIA refines each spot position through an iterative Gaussian-mask centroid algorithm running up to 300 iterations per spot, whereas TrackMate uses a single-pass parabolic interpolation around the LoG response peak. The iterative refinement and per-spot PSF width estimation incurs additional costs but yield the sub-pixel localisation precision responsible for MaCADEMIA’s RMSE performance.

### MaCADEMIA Stoichiometry

Determining protein stoichiometry, the number of subunits in a complex, is critical for understanding cellular machinery. Single-molecule fluorescence microscopy enables this by directly counting subunits in individual complexes. The main techniques translate a complex’s fluorescence intensity into a subunit number, either by counting discrete photobleaching steps^7,43^ or by comparing its total brightness to a calibrated single-molecule standard ^8^. New approaches including Bayesian^44^ and machine learning^45^ based photobleach counting are exciting but are not yet applicable to a wide range of stoichiometries. Other methods like Number & Brightness (NCB) are sensitive to autofluorescence and require precise calibration ^47^ while all intensity based approaches require that the brightness standard be measured under conditions matching those of the protein complex. Complete labelling of all proteins of interest is essential for accurate quantification of stoichiometry so endogenous genetic fusion with fluorescent proteins is often used and fluorescent protein maturation must also be taken into account. ^46^

MaCADEMIA determines protein stoichiometry by quantifying the step-wise photobleaching of fluorescently tagged complexes from timelapse images^48^. When imaged with high-power excitation light, individual fluorophores photobleach stochastically over time. The software tracks each complex and analyses its intensity trace, which often resembles a staircase as each subunit’s fluorophore photobleaches. This characteristic allows for the direct counting of subunits within the complex. This entire process can be performed on multiple colour channels simultaneously, enabling the analysis of multi-protein assemblies.

The analysis pipeline begins with calibration to determine the characteristic intensity of a single fluorophore. To do this, the intensity distribution of all detected molecules in a monomeric reference sample is measured with the peak of this distribution representing the characteristic brightness of a single fluorophore. To calibrate the single-molecule imaging platform, we characterised the photophysical properties of surface-immobilised monomeric mGFP. Analysis of 12,901 individual fluorescent spots revealed a unimodal intensity distribution with a peak at 596 ± 156 camera detector counts (in arbitrary units A.U.), defining the characteristic brightness of a single eGFP molecule (I_single_) on our system (Figure 3b). The photobleaching lifetimes of the population revealed a linear decay profile on a semi-logarithmic scale. A weighted least-squares fit to the distribution yielded a characteristic lifetime (τ) of 0.008 seconds, consistent with a first-order stochastic process (Figure 3c). Representative intensity-time traces are shown in (Figure 3d and Supplementary figure 4). Traces display stable intensity plateaus consistent with I_single_ before undergoing a single, photobleaching step to the background level. Application of the Chung-Kennedy edge-preserving filter (red line) preserved the step structure while reducing noise. These results confirm the successful immobilisation of monomeric eGFP and provide calibrated values for single-molecule brightness and photostability, which serve as a fundamental ruler for subsequent stoichiometry experiments.

**Figure 3:**
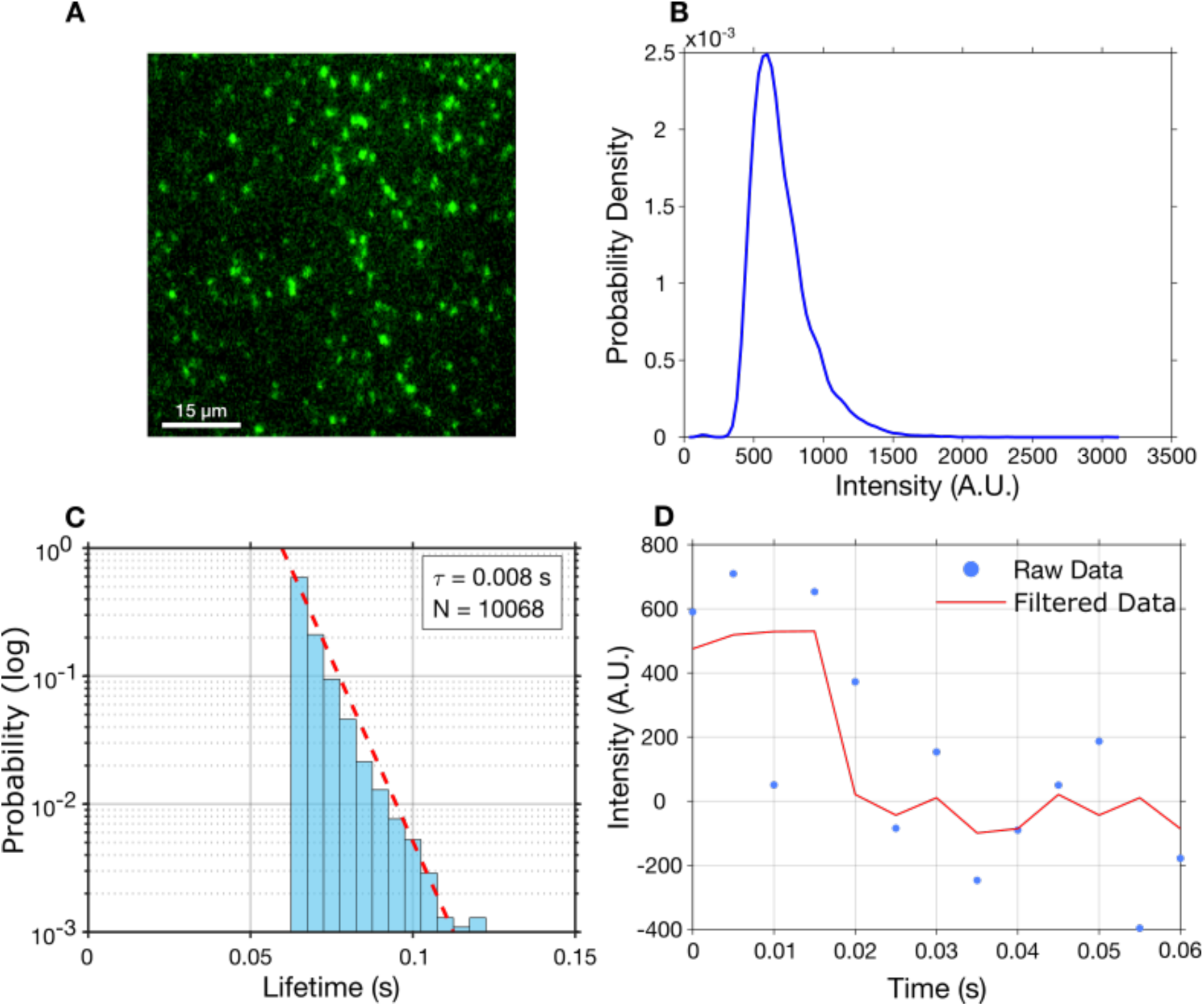
Photophysical characterisation of surface-immobilised monomeric eGFP for stoichiometry calibration. **(A)** Fluorescent image of monomeric GFP bound to a coverslip and imaged in HILO**. (B)** Probability density distribution of initial ffuorescence intensities from 12,S01 individual trajectories. The peak at 5SC A.U. defines the characteristic brightness of a single eGFP molecule (I_single_). **(C)** Semi-log distribution of photobleaching lifetimes. The linear decay indicates single-exponential kinetics consistent with a first-order stochastic process. The red dashed line represents a weighted least-squares fit yielding a characteristic lifetime (τ) of 0.008s. **(D)** Representative intensity-time trace of a single eGFP molecule (blue points). The trace exhibits a stable plateau at I_single_ followed by a single photobleaching step. The red line indicates the trace after Chung-Kennedy edge-preserving filtering.

To validate the stoichiometry analysis pipeline, the Nuclear Pore Complex (NPC) in U2OS cells with endogenously tagged Nup96-GFP was used as a biological standard.^49^ Each NPC contains exactly 32 copies of the Nup96 protein, providing a ground truth for stoichiometric quantification (Figure 4A). Ten individual cells were imaged with single-molecule HILO microscopy, each presenting a high density of NPCs distributed across the nuclear envelope, (Figure 4B). Following the analysis, the compiled stoichiometry data revealed a single, prominent population with a distribution mode of 31.3 ± 1.4 SEM (Figure 4C). This experimentally derived value is consistent with the theoretical stoichiometry of 32. The result, therefore, supports the method’s capability for providing accurate stoichiometric quantifications of large protein complexes *in situ*.

**Figure 4:**
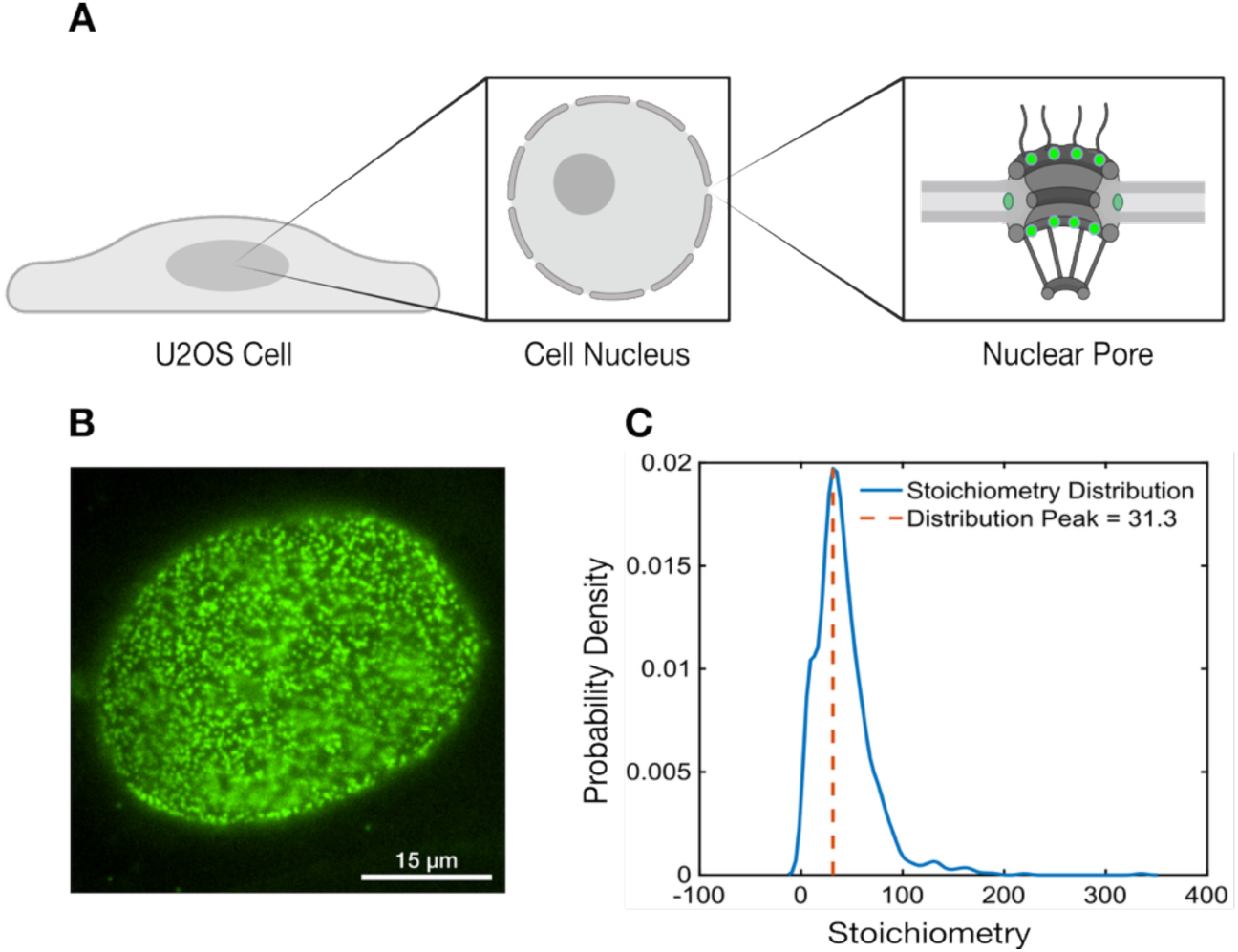
Quantification of Nuclear Pore Complex (NPC) stoichiometry. **(A)** Schematic illustration defining the hierarchy of the analysis, from the whole cell to the nuclear envelope region of interest (ROI), and finally the individual NPC structure. Green highlights indicate the target ffuorescent proteins. **(B)** Fluorescence microscopy image of a nucleus expressing NupSC-GFP. Distinct puncta correspond to individual NPCs on the nuclear envelope. **(C)** Probability density distribution of the calculated stoichiometry per NPC. The main peak is centred at 31.3, consistent with the expected stoichiometry of the complex (32).

### MaCADEMIA Simulation

To provide an essential ground truth for benchmarking the performance of image analysis algorithms, MaCADEMIA includes a simulation suite. MaCADEMIA allows the user to generate artificial datasets where every parameter is defined, it enables the user to systematically test analysis methods without the ambiguities inherent in experimental data.

The robust simulation pipeline allows the user to accurately model the key components of a real microscopy image from the ground up. The user can define idealised objects, or spots, with specific positions and intensities, program various types of motion like Brownian diffusion to create a known set of trajectories and finally add noise to simulate camera readout and other noise. To create the final realistic image, the simulation package accounts for the microscope’s optics by convolving each object’s position with the microscope’s Point Spread Function (PSF). For extended 3D objects, we also provide a more complex approach that integrates the PSF over the volume of the object’s defined shape. (Supplementary figure 5)

To demonstrate the utility of our simulation algorithms, we used them to generate ground truth data to test different sets of tracking parameters used in past imaging experiments with different foci densities (P_Wide, P_Medium and P_Narrow for low, medium and high foci density). Simulated images with spots of known intensities and positions were generated using a typical single-molecule microscope PSF (λ = 488 nm, NA = 1.45, FWHM ≈ 172 nm), which remained constant across all simulations. These were tracked using varied SNR thresholds and parameter configurations. Three parameter configurations were compared using different spot region of interest sizes (Fig. 5 and table 1).

**Figure 5:**
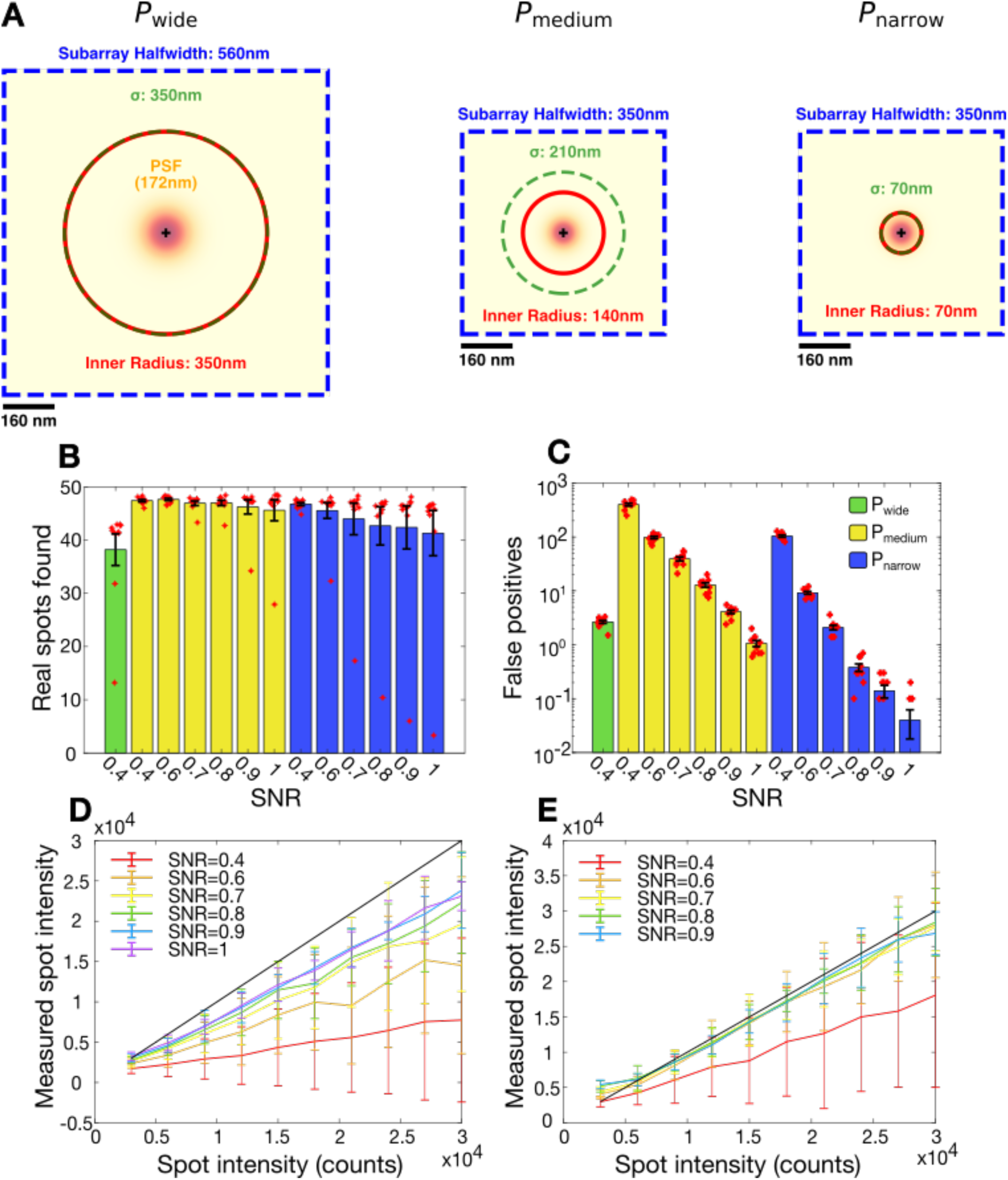
Validation of spot detection parameters using the simulation suite. Benchmarking of detection efficiency and photometric accuracy using simulated images with known ground truth. **(A)** Schematic showing three parameter configurations for spot detection analysis. The subarray halfwidth (blue box), inner integration radius (red circle), and Gaussian smoothing mask σ (green circle) vary across parameter sets whilst the PSF (172 nm FWHM) remains constant. Scale bars: 1C0 nm. **(B)** The number of real spots found (True Positives) across SNR ranges 0.4–1 for three parameter sets (P0,P1,P2). Suboptimal parameters (P1) captured significantly fewer spots than optimised sets (P2) (p=0.0072). **(C)** Quantification of False Positives (spurious detections) for the tested parameter configurations. **(D-E)** Photometric accuracy validation showing the correlation between measured spot intensity and ground-truth spot intensity. The data closely adhere to the line of identity (black line, y=x), confirming accurate intensity quantification.

**Table 1:** Localisation parameter sets evaluated in the tracking simulations. Three configurations (P_Wide, P_Medium, P_Narrow) varied the subarray half-width, inner circle radius and Gaussian mask sigma. Values are radii given in nanometres and as multiples of the simulated PSF FWHM (172 nm; λ = 488 nm, NA = 1.45, a diameter measurement). All other simulation parameters were held constant.

| Parameter Set | Subarray Halfwidth | Inner Circle Radius | Gauss Mask Sigma |
|---|---|---|---|
| P_Wide (P0) | 560 nm, 3.3× FWHM | 350 nm, 2.0× FWHM | 140 nm, 0.8× FWHM |
| P_Medium (P1) | 350 nm, 2.0× FWHM | 140 nm, 0.8× FWHM | 70 nm, 0.4× FWHM |
| P_Narrow (P2) | 350 nm, 2.0× FWHM | 210 nm, 1.2× FWHM | 70 nm, 0.4× FWHM |

Analysis revealed that parameter selection involves important tradeoffs. Comparing P_wide (P0) with P_medium (P1) showed that reducing spatial integration parameters (inner circle radius, Gaussian mask sigma) decreased detection efficiency, with P_medium capturing significantly fewer real spots (Figure 5A, p=0.0072). This reflects a tradeoff: narrower integration parameters reduce localisation precision by integrating fewer photons per spot. However, this precision loss can be compensated by increasing the SNR detection threshold (Figure 5B), which rejects lower-quality spots whilst controlling false positive rates across all parameter sets. The SNR threshold proved critical for accurate intensity quantification. An SNR threshold of 0.4 substantially underestimated spot intensities, with measured values falling well below the line of identity (Figure 5C,D). In contrast, higher thresholds (SNR ≥0.8) yielded accurate measurements following the expected 1:1 relationship between measured and ground-truth intensities. These results demonstrate that incorrect parameter selection introduces systematic measurement errors undetectable without ground-truth validation, and that MaCADEMIA’s simulation framework provides a principled approach for parameter optimisation.

### MaCADEMIA Diffusion

To determine the diffusion coefficient (*D*) of tracked particles, MaCADEMIA offers several analysis methods, primarily based on Mean Squared Displacement (MSD). The software calculates the average squared distance a particle travels over increasing time lags (τ). For particles exhibiting simple Brownian motion, a linear plot of MSD versus τ is generated, and the diffusion coefficient is extracted from the slope according to the relation MSD(τ)=2d*D*τ, where *d* is the dimensionality of the motion.

For more complex analysis in heterogeneous environments (Figure 6A), the software can calculate micro-diffusion coefficients to capture dynamic changes in mobility. This is achieved by calculating the MSD over a short, initial time window for each trajectory, providing a localised diffusion coefficient for each track.^50^ By pooling these values from all tracks, a distribution of micro-diffusion coefficients is generated, a powerful output that can reveal distinct subpopulations with different mobilities (e.g., mobile vs. confined).

**Figure 6:**
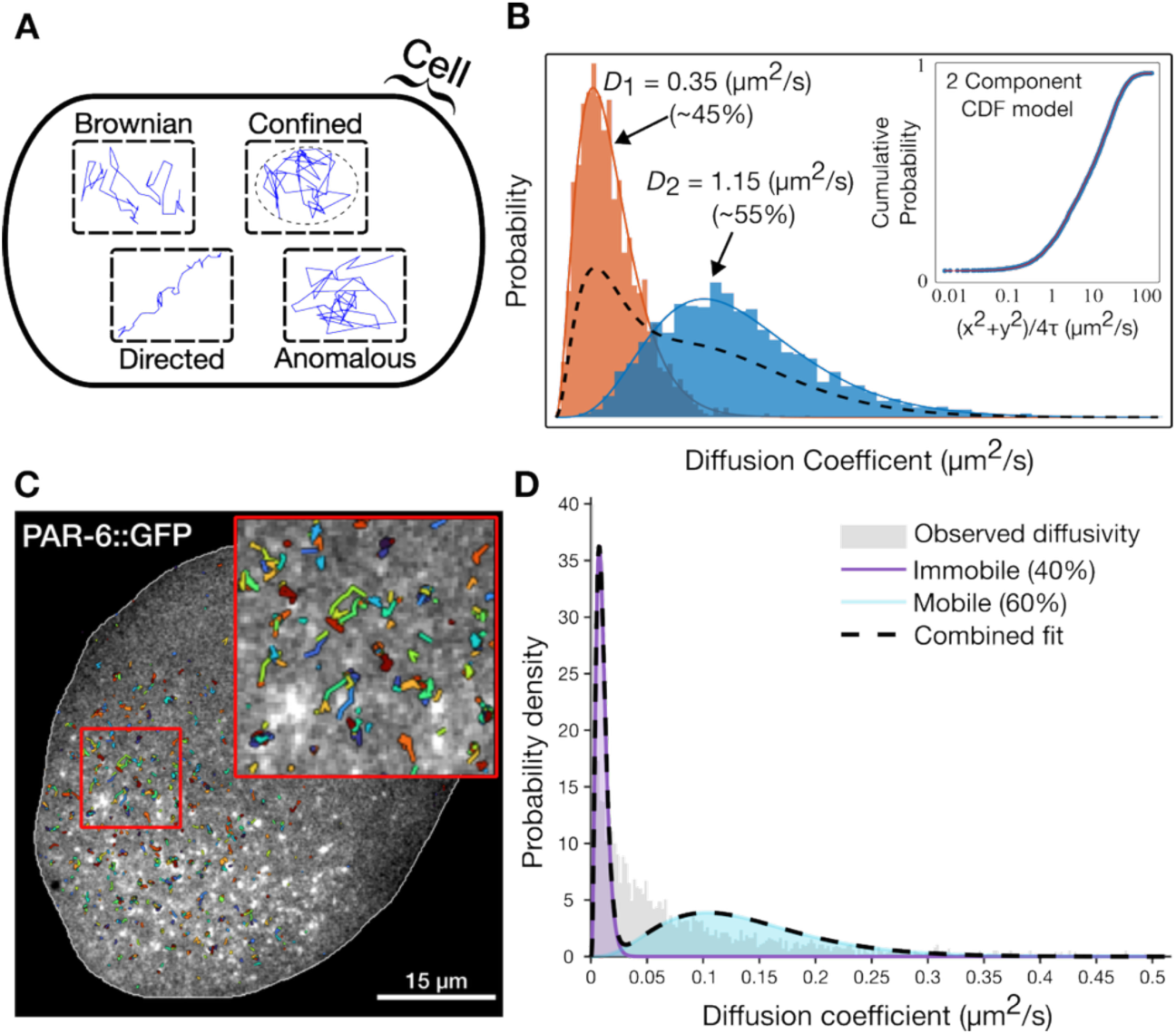
Diffusion analysis and mobility modelling. **(A)** Schematic representation of different particle motion modes within a cell: Brownian, Confined, Directed, and Anomalous diffusion. **(B)** Distribution of diffusion coefficients (D) derived from single-particle trajectories. The histogram is fitted with a 2-component model, identifying distinct populations with different mobilities (e.g., D1≈0.35μm2/s and D2≈1.15μm2/s). The inset shows the Cumulative Distribution Function (CDF) fit used to validate the model. **(C)** Representative ffuorescence image of a masked C. elegans embryo overlaid with single-particle PAR-C trajectories generated by MaCADEMIA. For visual clarity, only tracks persisting for longer than 5 frames are displayed. Inset shows zoomed ROI. **(D)** Probability density of pooled PAR-6 apparent diffusion coefficients (D). A two-component MLE fit (black dashed line) resolves the data into immobile (purple, 40%) and mobile (cyan, C0%) populations

To validate the suitability of the statistical model used by MaCADEMIA, we performed Monte Carlo simulations of theoretical diffusion distributions. We generated synthetic datasets containing known mixtures of distinct mobility states (e.g., 45% slow, 55% fast). Fitting these synthetic distributions with the multi-component Gamma model employed by our pipeline successfully recovered the underlying parameters (Figure 6B). This confirms that our chosen statistical approach can accurately quantify the diffusion characteristics of heterogeneous populations (*D*_1_ ≈ 0.35*μm*^2^/*s*, *D*_2_ ≈ 1.15*μm*^2^/*s*).

Finally, as a robust alternative to standard MSD, we offer analysis based on the cumulative distribution functions (CDFs) of particle displacements.^4^ This approach is less sensitive to trajectory length and is particularly effective for modelling systems with multiple mobility states, providing a complementary method for validating diffusion measurements. As shown in the inset of Figure 6B, the 2-component CDF model provided a precise fit to the simulated data, offering a complementary method for validating diffusion measurements.

To validate MaCADEMIA’s diffusion analysis using real biological data, we analysed the diffusion behaviour of the polarity protein PAR-6, which has previously been shown to exhibit at least two distinct mobility states at the cortex of *C. elegans* zygotes ^51^. Embryos were imaged using single-molecule near TIRF microscopy (Figure 6C). MaCADEMIA autonomously partitioned the trajectories into a highly restricted, immobile fraction (40.2%, D=0.0099 μm^2^/s) and a distinct, freely diffusing mobile population (59.8%, *D*=0.1380 μm^2^/s). This unsupervised classification closely mirrors the previous observations for PAR-6, where a mobile fraction (∼43% at *D*=0.17±0.10 μm^2^/s) and immobile fraction (∼22% at *D*=0.008±0.008 μm^2^/s) were also observed. Ultimately, this demonstrates MaCADEMIA’s capacity to extract accurate, biophysically relevant mobility states from dense tracking data without relying on user-defined cut-offs.

### MaCADEMIA Colocalisation

Dual colour single-molecule microscopy allows protein-protein and other molecular interactions to be measured. This is achieved by quantifying the spatial overlap of the PSFs of distinct molecular species over time. MaCADEMIA implements this in a rigorous co-localisation workflow. The process begins with computational image registration to correct for chromatic aberrations and spatial shifts, ensuring that pixel coordinates across channels correspond to identical physical locations. Without this alignment, derived spatial correlations would be inaccurate.

While pixel-based methods typically rely on global intensity correlation, MaCADEMIA employs a discrete, object-based approach optimised for single-molecule data. This is demonstrated here using the polarity proteins PAR-3 (Channel 1) and PAR-6 (Channel 2), which have been shown *in vivo* to form part of the same protein complex in *C. elegans* embryos ^52^. Emitters are detected and tracked independently within each channel (Figure 7A-B) before being computationally linked. Inter-channel association is governed by strict spatial constraints: a PAR-3 trajectory is assigned to a PAR-6 partner only if the pair falls within a user-defined search radius and exceeds a minimum Point Spread Function (PSF) overlap integral threshold.

**Figure 7:**
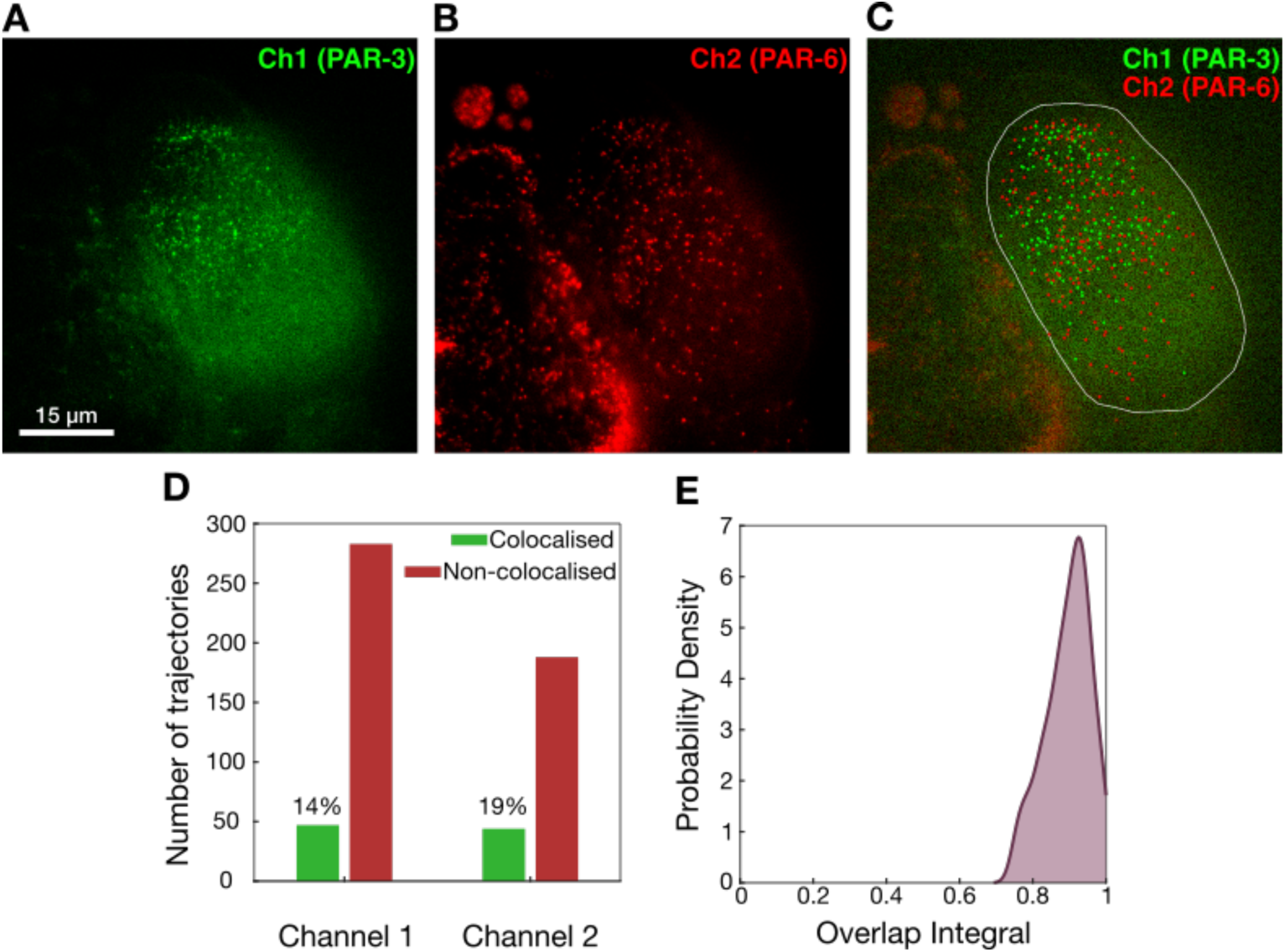
Object-based colocalisation analysis of PAR-3 and PAR-C. **(A-B)** Representative single-molecule microscopy images of **PAR-3** (Channel 1) and **PAR-C** (Channel 2). **(C)** Composite overlay showing detected particle trajectories (**PAR-3: Green**, **PAR-C: Red**). MaCADEMIA performs independent tracking in each channel before computationally linking trajectories based on strict spatial proximity and Point Spread Function (PSF) overlap criteria. ***(D)*** *Quantification of linkage rate, displaying the proportion of trajectories for each protein classified as colocalised versus independent in each channel. **(E)** Distribution of the PSF overlap integral for linked pairs. High integral values confirm genuine molecular overlap, distinguishing true colocalisation from stochastic proximity*.

This workflow visualises valid interactions directly on the composite overlay (Figure 7C) and quantifies linkage efficiency to distinguish between colocalised and independent populations (Figure 7D). Furthermore, the spatial fidelity of these interactions is validated by analysing the distribution of overlap integrals (Figure 7E). MaCADEMIA found an average of ∼16% of trajectories were colocalised across the two channels consistent with previously reported estimates ^52^.

### MaCADEMIA Segmentation

Traditional algorithms like intensity thresholding (e.g Otsu’s method ^41^), the watershed transformation, and edge detection form the foundation of image segmentation. These methods are sensitive to noise and poor contrast. Recently Artificial Intelligence (AI) based methods have been developed with high levels of general performance^42^. Deep learning models, such as Convolutional Neural Networks (CNNs) ^55^, excel by learning complex features directly from data, achieving high accuracy even in challenging images. Despite the power of AI, traditional techniques remain crucial in scientific analysis. Unlike many AI black-boxes, they are transparent, computationally inexpensive, and do not require large, annotated training datasets. For images with well-defined features, traditional algorithms are often more practical and efficient. The choice of method, therefore, depends on the complexity of the imaging problem, not the novelty of the method itself.

To accurately identify and isolate objects of interest, such as individual cells or organelles, MaCADEMIA provides a powerful and tuneable workflow for image segmentation. This process is particularly critical for subsequent analyses like optical proteomics, where the absolute protein copy number is calculated from the total fluorescence within a precisely defined volume. Here, the actual physical boundary of the cell must be determined for accurate copy number determination.^46,56^ For this reason, we prioritise methods that are transparent, deterministic, and fully interpretable over black-box AI/ML approaches.

Our algorithmic workflow is efficient, requires no training data, and involves logical steps that can be customised to suit specific data. After optional pre-processing steps to reduce image noise, we provide a suite of core techniques. These include intensity thresholding to create a binary mask based on pixel brightness, edge detection to identify object boundaries, and the powerful watershed algorithm, which treats the image like a topographical map to separate touching or clustered objects. The true strength of our module is the synergy of these tools; a sequential and fully auditable workflow for accurate segmentation, ensuring the reliability of downstream quantitative measurements.

Cellpose is an open source, deep learning-based tool for cell and nucleus segmentation. ^20^ Cellpose3.0 offers the generalist “cyto3” segmentation model, trained off ∼8,402 images of diverse cell types and imaging modalities from 9 combined datasets. A MaCADEMIA adaptive Otsu thresholding model with watershedding was tuned to a set of fixed microglia cells (HMC3) stained with Wheat Germ Agglutinin (WGA) Alexa Fluor™ 488 conjugate. This thresholding model, available in MACADEMIA, was compared to the cyto3 generalist model (Figure 8) run on both the WGA and oblique channels. The adaptive Otsu with watershedding performs similarly to Cyto3 Cellpose on the WGA channel in terms of Jaccard Index similarity and normalised false positives and identified no missed cells. The cyto3 generalist model performed significantly worse on the oblique channel; however, a Cellpose model trained specifically on microglia achieved comparable performance to the adaptive Otsu method; though such trained models require substantial annotated training data and remain less interpretable than traditional algorithmic approaches.

**Figure 8:**
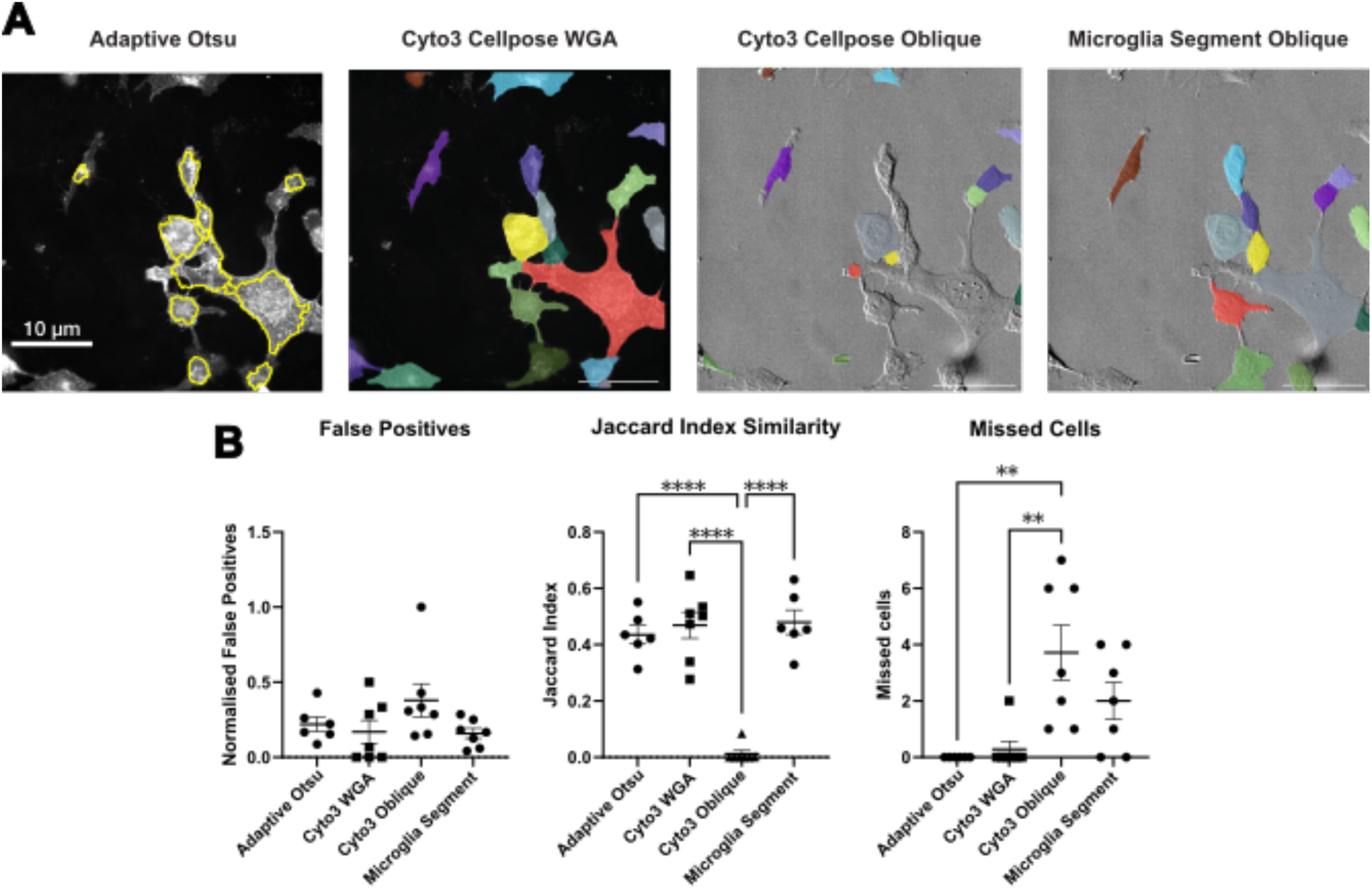
Comparison of segmentation methods for fixed HMC3 microglia cells. **(A)** Exemplar masks for the methods tested. **(B)** Quantitative comparison of segmentation performance across methods.

### MaCADEMIA Optical proteomics

Where MaCADEMIA stoichiometry functions determine the number of molecules in a single complex, optical proteomics determines the absolute protein copy number within a whole cell or sub-cellular organelle. Optical proteomics deconvolves the total fluorescence intensity within fluorescence images to determine the underlying number of molecules generating that intensity.^57^ Theoretically the copy number of fluorescent molecules in a cell could be determined by summing the intensity within the cell and dividing it by the characteristic intensity of a single fluorophore (Isingle). However, for large enough objects the depth of field of the microscope must be factored in, as some fluorescence from emitters far from the focal plane may appear out with the object. And for compartments within cells (e.g. the nucleus (Fig. 9)), the contribution of fluorescence from other compartments (e.g. the cytoplasm) must also be controlled for.

**Figure 9:**
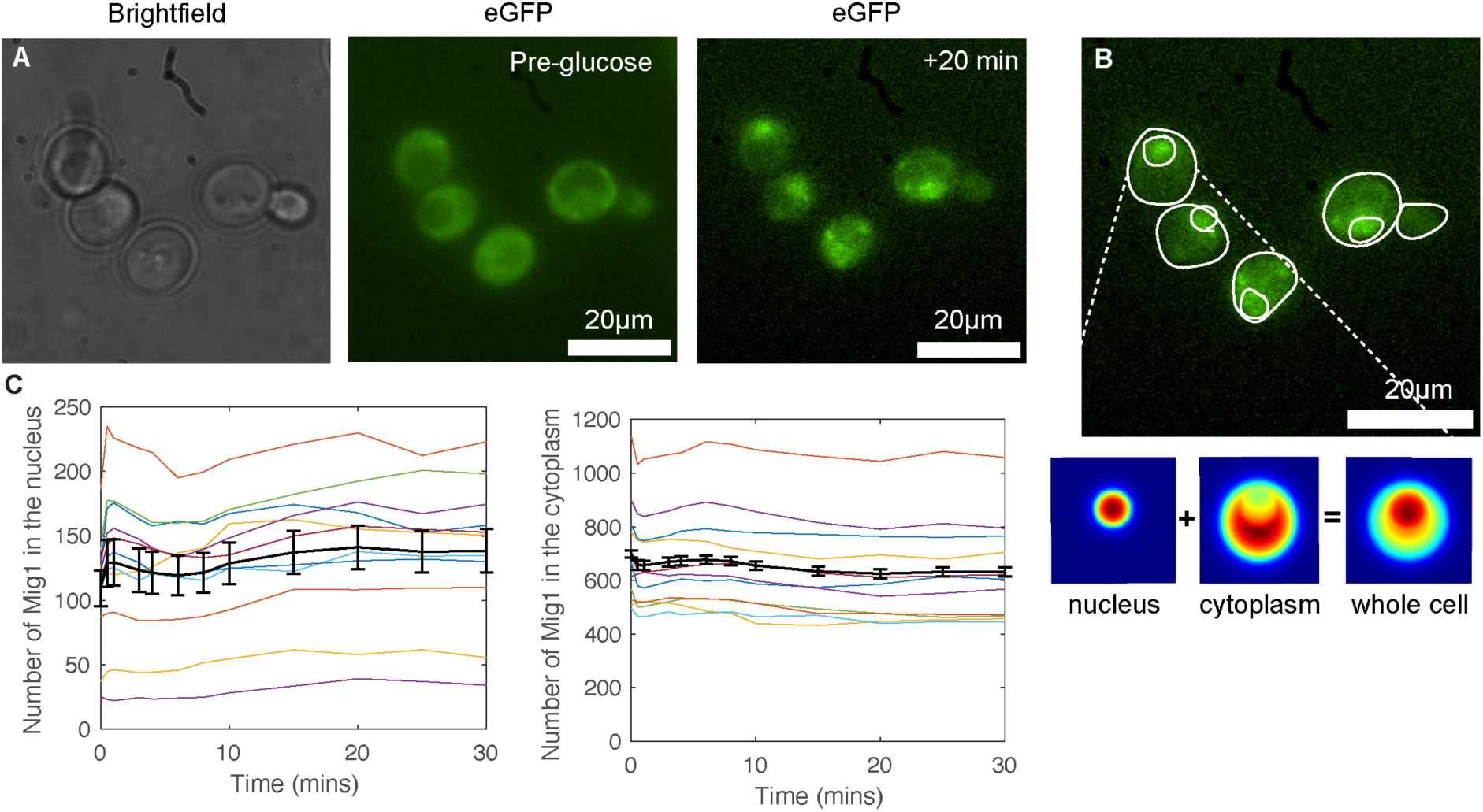
A. Micrographs of S. Cerevisiae expressing a GFP tagged glucose responsive transcription factor, Mig1, before and after exposure to extracellular glucose. B. Schematic illustrating MaCADEMIA for optical proteomics. Cells are segmented for the cell and nucleus (white outlines), before a model PSF is integrated over the compartment volumes to create model volumes which can be used to deconvolute the original image. C. the total number of Mig1 in the nucleus and cytoplasm as a function of time.

To determine the absolute protein copy number in distinct cellular regions, MaCADEMIA employs a workflow that models the three-dimensional nature of the cell. The process begins with the 2D segmentation masks of the cell and any internal compartments, such as the nucleus.

To account for fluorescence far from the focal plane or from other compartments, the algorithm generates a 3D projection model based on the cell’s geometry. The software is flexible and can construct this model using different shapes to match the organism being studied; for instance, by assuming a spherical shape for mammalian cell nuclei or a rod shape for bacteria. It then simulates theoretical images based on these shapes and the system’s point spread function (PSF). These simulated images represent how a uniform concentration of protein in each compartment would appear after being imaged.

The core of the analysis is a multiple linear regression model.^56^ To avoid bias from bright, oligomeric spots, these regions are masked out from the experimental image. The model then relates the measured pixel intensities in the remaining “pool” regions to the corresponding intensities in the simulated compartment projection images. By solving this relationship, the model accurately deconvolves the overlapping signals and determines the distinct protein concentration for each compartment.

Finally, these concentrations are multiplied by the respective compartment volumes (calculated from the geometric model) to yield the total number of molecules in each pool. The software also quantifies molecules within bright spots and applies a 3D correction, providing a comprehensive budget of the protein’s cellular distribution.

MaCADEMIA optical proteomics was demonstrated by deconvolving the fluorescence signal from a GFP tagged glucose responsive transcription factor, Mig1, in S. cerevisiae, in response to a glucose shift. (Fig. 9). Mig1 translocates into the nucleus in response to glucose (Fig. 9C).

## Discussion

The analysis of single-molecule localisation microscopy data has long been hampered by a fragmented and highly specialised software landscape. MaCADEMIA directly addresses this critical gap by providing a single, integrated, and comprehensive MATLAB-based platform. It consolidates the entire analytical pipeline, from essential pre-processing steps like cell segmentation to quantitative measurements of protein stoichiometry, diffusion, colocalisation, and absolute protein copy number into one package.

A core strength of MaCADEMIA lies in its commitment to algorithmic transparency, setting it apart from opaque black-box machine learning approaches. This design philosophy ensures that every step of the analysis is interpretable, tuneable, and scientifically auditable, giving researchers full control over their experimental pipeline and confidence in the results. Furthermore, a powerful simulation suite provides an essential tool for validation and methods development, allowing users to benchmark the software’s performance against a known ground truth.

By consolidating these tools into a cohesive, user-friendly environment, MaCADEMIA significantly lowers the technical barrier for researchers aiming to extract profound quantitative insights from live-cell imaging data. It allows biologists to move beyond qualitative observation to robustly measure the dynamics, architecture, and interactions of molecular complexes within their cellular context. Ultimately, MaCADEMIA stands as a powerful and accessible toolkit poised to enhance the reproducibility and depth of discovery in the expanding field of quantitative cell biology.

## Code availability

MaCADEMIA requires MATLAB 2021b or later and the statistics & machine learning and image processing toolboxes. MaCADEMIA is open-source and available at https://github.com/awollman/MaCADEMIA.

## Data availability

The data collected and analysed in this work is available at http://doi.org/10.5281/zenodo.23361879.

## Acknowledgements

This work was supported by funding from the Academy of Medical Science (springboard award, SBF007\100046) and Royal Society Paul Instrument Fund (Fund PIF\R1\261033) to AW, EPSRC to ML (EP/Y000501/1) and the BBSRC responsive mode to JR (BB/R019436/1).

## Methods

### GFP purification

GFP was overexpressed in E. coli BL21 (DE3) cells using pET28 vectors. Cultures were grown to OD₆₀₀ ∼0.6, induced with 0.5 mM IPTG, and incubated overnight at 18°C. Cells were harvested by centrifugation, washed with PBS, and stored at −80°C.

Pellets were resuspended in lysis buffer (50 mM HEPES, 200 mM NaCl, pH 7.5) containing lysozyme (100 μg/mL), DNase I (10 μg/mL), and protease inhibitor, then lysed using an Avestin Emulsiflex homogenizer at 2-3 bar. Following ultracentrifugation and filtration, soluble protein was purified using a 5 mL HisTrap HP column on an ÄKTA Start system with imidazole gradient elution. GFP-containing fractions were identified by colour and SDS-PAGE, dialyzed overnight against storage buffer, flash frozen, and stored at −80°C.

### GFP imaging

To calibrate single-fluorophore brightness and validate stepwise photobleaching analysis, immobilised purified GFP was used as a monomeric reference. Plasma-cleaned coverslips were coated with anti-GFP antibody (Sigma; G6539, 1.5 μg/ml), then incubated with purified GFP (1 μg/ml) before unbound protein was removed by washing with PBS. Individual GFP molecules were thereby spatially isolated and predominantly monomeric, providing a well-defined single-fluorophore reference. Immobilised GFP was imaged under identical conditions to experimental samples until complete photobleaching. The intensity distribution of detected spots was taken to represent single-GFP brightness under these imaging conditions and was used to determine fluorophore-to-protein stoichiometry in experimental datasets.

### HMC3 cell staining and imaging

5K HMC3 cells per well were seeded in PhenoPlate 96-well Cyclic Olefin bottomed black walled plates (Revvity) in high glucose supplemented DMEM. 24 hours later, cells were loaded with 500nM MitoTracker™ Orange CM-H2TMRos (ThermoFisher), reconstituted in anhydrous DMSO (ThermoFisher), by 30min incubation (37 °C,5% CO2) in serum free supplemented high glucose DMEM for 30mins, washed once with DPBS, then treated with a range of benchmarking stimuli, singly or in combination, supplemented DMEM under normal culture conditions. Post-treatment, cells were washed once with DPBS, fixed in 4% formaldehyde (Thermofisher), counterstained with 3uM Hoechst 33342 (Thermofisher) for 15 minutes, then incubated with 1ug/mL Wheat Germ Agglutinin conjugated to Alexa Fluor 488 (WGA-A488; Thermofisher) in DPBS for 10minutes, both incubated at room temperature protected from light. After staining, cells were washed 3 times with DPBS. Images were acquired at 20x magnification on the Zeiss Cell Discoverer 7 widefield automated high content imaging microscope equipped with a 7 LED Colibri light source (365, 410, 470, 511, 567, 590, 625nm LED’s) using Filterset 90 HE LEDusing a Plan-Apochromat 20x/0.7 NA objective. WGA-A488 was detected at 50ms exposure time 2.11uM depth of focus and MitoTracker™ Orange CM-H2TMRos at 250ms 2.35uM depth of focus at 100% light source intensity, Hoechst33342 at 20.0% light intensity 25ms exposure time and 1.86um depth of focus, and transmitted light (TL) oblique at 10.0% light intensity 2ms exposure time 2.25uM depth of focus. Images were acquired with a CMOS camera (Hamamatsu Fusion).

### U2OS Culture and imaging

U2OS CRISPR NUP96-mEGFP cells (300174, Cytion) were cultured in phenol red free DMEM (21063029, Thermofisher) supplemented with 10% FBS and Pen/strep. All cells were maintained at 37C with 5% CO2. 1 day before imaging cells were seeded into ibidi 2 well slides in the same phenol red free DMEM. Cells were imaged the following day.

### C. elegans culture and imaging

*C. elegans* strains used in this study were maintained on standard nematode growth media (NGM) under routine laboratory conditions ^58^. PAR-6 diffusivity was measured in strain JJ1579 (unc-119(ed3) III; zuIs77 [par-6p::PAR-6::GFP; unc-119(+)] (Figure 6). Dual-colour imaging was performed with strain NWG0198 (par-3(it298 [par-3::gfp]) III; par-6(cp346[PAR-6::HaloTag]) I).

For imaging embryos were dissected in 3 μl of egg buffer (2 mM CaCl₂, 118 mM NaCl, 48 mM KCl, 2 mM MgCl₂, 25 mM HEPES, pH 7.4) on a 30 mm circular coverslip (#10343435, Fisher Scientific). The coverslip was then inverted onto a 2% agar pad made with the same egg buffer and placed in a custom-made temperature-controlled stage holder (Bioptechs Oasis Cooling System). To improve thermal stability (20 °C), the objective lens was temperature-regulated using an objective cooling collar (Bioptechs). TIRF imaging was performed on a bespoke single-molecule TIRF microscope constructed around a Nikon Ti-E microscope body, using an Obis 488 laser set to 20 mW, beam expanded to ∼20 µm and using highly inclined illumination (HILO). Imaging was performed using Photometrics Evolve 512 at 100 nm/pixel with a 100x Nikon 1.49NA TIRF objective lens. PAR-6::GFP and PAR-6::HaloTag/PAR-3::GFP were imaged at the anterior cortex of mixed stage zygotes, subarraying the camera sensor to a ROI of 128×128 pixels to achieve 15ms/frame exposure time and 5s acquisitions intervals.

For live imaging of HaloTag fusion proteins, adult worms were cultured overnight in S-medium (150 mM NaCl, 1 g/L K₂HPO₄, 6 g/L KH₂PO₄, 5 mg/L cholesterol, 10 mM potassium citrate pH 6.0, 3 mM CaCl₂, 3 mM MgSO4, 65 μM EDTA, 25 μM FeSO₄, 10 μM MnCl₂, 10 μM ZnSO₄, 1 μM CuSO₄, 1 mM IPTG, 10 μg/ml carbenicillin, 10 μg/ml tetracycline, and 100 U/ml nystatin) supplemented with JFX549 HaloTag ligand (1.25 μM, Janelia Farm) at 15 °C under gentle rotation (30–50 worms in 65 μl of culture in a 0.5 ml tube). After incubation, worms were washed six times in M9 buffer with 0.01% Triton X-100 (M9: 22 mM KH₂PO₄, 42 mM Na₂HPO₄, 86 mM NaCl, 1 mM MgSO₄) to remove excess dye and bacteria and transferred to fresh NGM plates until imaging.

### Yeast culture and imaging

*S. cerevisiae* yeast expressing genomically tagged Mig1::GFP was grown in YPD medium with 4% glucose at 30 °C, 180 rpm. Overnight cultures were subcultured to OD₆₀₀ 0.1 and grown to OD₆₀₀ 0.6. Cells were pelleted, washed 3× with YNB-glucose, resuspended in YNB-glucose, and transferred to Concanavalin A-coated ibidi μ-Dishes for imaging.

Imaging was performed on a Zeiss 880 Confocal microscope using a Plan Apo 63X/1.4 oil objective at 30 °C. Cells were imaged in the glucose-starved state, then at 5-minute intervals for 30 minutes following addition of 4% glucose.

### Single particle tracking

First, the software enhances potential fluorescent signals by applying a Laplacian of Gaussian (LoG) filter, which highlights spot-like features of a specified radius while suppressing broad background variations. To separate these features from noise, an adaptive intensity threshold is automatically calculated for each frame based on the image’s pixel statistics. Specifically, the threshold is determined by calculating the median and standard deviation of the background pixels, ensuring the segmentation is statistically robust to fluctuations in noise levels throughout the movie. The resulting filtered image is converted into a binary mask by identifying regional intensity maxima that exceed this threshold.

Next, this binary mask is refined to ensure only high-confidence candidates are processed. The software identifies potential spot centers and removes candidates located too close to the image borders to allow for accurate quantification. This process efficiently identifies the initial coordinates for all local intensity maxima across the field of view.

Finally, each candidate spot is precisely localized and quantified. A candidate detection is accepted only if its signal-to-noise ratio (SNR) exceeds a minimum threshold (default SNR_min = 0.4). The software refines the spot’s position to sub-pixel accuracy using an iterative Gaussian-weighted centroid algorithm, which repeatedly calculates the center of mass until the coordinates converge. The spot’s characteristic intensity is measured by summing the pixel values within a circular region (default inner_circle_radius = 5 pixels), which is large enough to encompass the microscope’s point spread function (PSF). To ensure accuracy, the local background is calculated from pixels outside this signal mask and subtracted from the total integrated intensity. While these parameters are optimised defaults, they are fully user-adjustable to accommodate specific optical setups. Each pipeline stage offers algorithm options: LoG or morphological detection, iterative or quadratic localisation, and LAP (with gap closing) or greedy linking. Defaults prioritise accuracy while alternatives offer compatibility or speed.

Detection accuracy was measured using the Jaccard Index (J), calculated as:

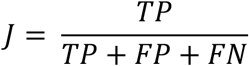

Where TP is the number of true positives, FP is the number of false positives, and FN is the number of false negatives. Localisation accuracy was measured by the Root Mean Square Error (RMSE) in pixels.

Timing benchmarks were performed on a standard laptop (Apple M1 Pro, 16 GB RAM, macOS Sonoma 14.6.1) using the 100-frame, 512×512 pixel “vesicle high density SNR 7” sequence from the particle tracking benchmark of Chenouard et al. ^42^. MaCADEMIA runtime was measured in MATLAB R2024b using tic/toc around the full tracker_upgraded call with default parameters. TrackMate (version 7.14.0) was run in Fiji (2.16.0) via a Jython script with the LoG detector (radius 2.5 pixels, quality threshold 8.0, sub-pixel localisation enabled) and LAP tracker (max linking distance 5 pixels, max gap-closing distance 2 pixels, max frame gap 1). The TrackMate quality threshold was tuned to give a per-frame spot count of the same order of magnitude as MaCADEMIA on the same sequence. Each pipeline used five timed runs from which the mean was reported. Detection and tracking stages were timed separately for TrackMate; MaCADEMIA’s timing represents the combined pipeline.

### Stoichiometry

The stoichiometry, or the number of fluorescent subunits, for each diffraction-limited spot was determined by analysing its intensity-time trace. The analysis aimed to estimate the spot’s initial intensity *I*_0_ at the start of the observation period, which was then normalised by the characteristic intensity of a single fluorophore *I_Sin_*_g*le*_ to yield the subunit count.

To robustly estimate *I*_0_ and account for photobleaching, several methods were implemented. The most direct methods included using the intensity from the first recorded frame or calculating the mean intensity over a predefined number of initial frames. For traces exhibiting photobleaching, extrapolation techniques were employed. The initial portion of the intensity decay was fitted with a linear function *I*(*t*) = *mt* + *I*_0_ using a constrained linear least-squares fit, where the y-intercept represents the extrapolated intensity at time zero. Alternatively, the decay was modelled by fitting a single exponential function *I*_0_*e*^−*kt*^ + *C* using the non-linear least-squares algorithm from the MATLAB Curve Fitting Toolbox. This fit was performed either with a fixed, predetermined bleaching rate constant *k* or by allowing both the initial intensity *I*_0_ and the rate constant *k* to be optimised as free parameters.

### Simulation

The simulation package created time-lapse image series of a predefined number of particles undergoing two-dimensional Brownian motion within a defined area. In each frame, the displacement of each particle was determined by drawing from a Gaussian distribution with a mean of zero and a standard deviation of *σ* = √2*D*Δ*t* where *D* is the specified diffusion coefficient and Δ*t* is the frame time. The simulation also included an optional photobleaching routine, where in each frame, particles had a fixed probability of being removed and replaced by a new particle at a random location to mimic fluorophore burnout and replacement.

The ground-truth particle trajectories were then used to render a synthetic image stack. Each particle’s position in each frame was represented by a two-dimensional Gaussian profile, simulating the microscope’s point spread function (PSF). To replicate experimental noise sources, the resulting signal was first subjected to a Poisson noise model to simulate photon shot noise, after which Gaussian noise was added to the entire frame to simulate camera read-out noise and background fluorescence. The final synthetic image stack was saved as a multi-frame TIFF file and processed using the identical tracking and analysis software as the experimental data. The performance of the tracking algorithm was quantified by comparing the tracked spot positions against the known ground-truth coordinates, allowing for the calculation of false positive (spurious detections) and false negative (missed particles) rates.

### Diffusion

Diffusion analysis proceeded from calculating diffusion coefficients for individual trajectories to modelling the distribution of mobilities across the entire particle population. The apparent diffusion coefficient (*D*) for each individual trajectory was determined via Mean Squared Displacement (MSD) analysis. For each particle’s path, the time-averaged MSD was calculated for a series of time lags (*τ*). The initial, linear portion of the resulting MSD-versus-*τ* curve was then fitted to the theoretical relation for two-dimensional Brownian motion: *MSD*(*τ*) = 4*Dτ* + 4*σ*^2^ This fitting was performed using a weighted linear least-squares regression, where each MSD point was weighted by the number of individual displacements used in its calculation. The y-intercept of the fit, 4*σ*^2^, is related to the localisation precision of the measurement and was constrained during the fitting process. The diffusion coefficient *D* was subsequently determined from the slope of the fitted line.

This procedure yielded a distribution of diffusion coefficients from all valid trajectories. To identify and quantify distinct mobility states within this population, the empirical Cumulative Distribution Function (CDF) of the diffusion coefficients was constructed. This CDF was fitted using non-linear least-squares regression to models describing a mixture of one, two, or three mobile species, governed by the equation:

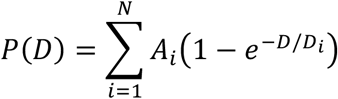

Where *N* is the number of components (1, 2, or 3), *A_i_* is the fractional population of each component (such that ∑ *A_i_* = 1) and is the characteristic diffusion coefficient of that component. The optimal number of components required to describe the experimental data was determined by comparing the reduced chi-squared (*χ*^2^) statistic for each model fit.

### Colocalisation

Spots detected in two separate channels were analysed for colocalisation. First, coordinates from the second channel were corrected with a pre-calibrated affine transformation to align the two imaging channels.

The colocalisation of individual spots was then determined. For a given pair of spots, an overlap integral was calculated based on the distance between the two spots and their respective point-spread function (PSF) widths. The specific formula used was:

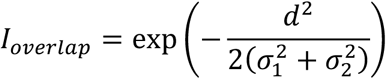

where d is the distance between the spot centres and *σ*_1_^2^and *σ*^2^ are the variances of their respective Gaussian PSFs. A spot in one channel was linked to a potential partner in the other channel if they were within a maximum search radius and their overlap integral (*I_overlap_*) was the highest among all candidates and exceeded a minimum threshold (e.g., 0.75). This assignment was constrained to be unique, ensuring each spot was paired with at most one partner. Depending on the experimental setup, this analysis was performed either on a frame-by-frame basis or between alternating frames for ALEX imaging.

Finally, colocalisation was assessed at the trajectory level. After performing single-particle tracking independently in each channel, the spot-linking algorithm was applied only to those spots belonging to valid trajectories. A trajectory in one channel was considered colocalised with a trajectory in the other if they contained linked spots. If a trajectory was linked to multiple partners over its lifetime, it was uniquely assigned to the partner trajectory with which it shared the highest number of colocalised frames. This process yielded pairs of linked trajectories, the duration of their interaction, and their mean separation distance.

### Cellpose segmentation and Jaccard Index calculation

Seven masks covering the full treatment range of the cells were generated on Cellpose3.0 using the Cyto3 model run separately on the WGA-A488 and Oblique channels with cell diameter set to 120 pixels. MATLAB code was used to compare the ground truth segmentation masks traced on the cellpose general user interface with the model generated segmentation masks to calculate the mean similarity as the Jaccard Index (meanSimilarity), number of missed cells (noMissedCells), and number of false positive (noFalsePositives). The mean Jaccard similarity reflects the degree of overlap between segmented cells and their corresponding ground truth cells. Missed cells represents the number of ground truth cells not matched by any segmented cell; false positives represent the number of segmented cells not matched by any ground truth cell. The MATLAB function works by identifying pixels belonging to each cell in the ground truth mask then looping through each cell in the model mask and if there is overlap by the element-wise product of the segmented single-cell matrices being greater than 0, the Jaccard similarity is calculated and inner loop of the code broken. No match occurs when the product is less than zero and a missed cell is computed. To calculate the false positives, a second similar loop compares the pixels representing cells in the respective model mask and loops through each cell in the ground state mask. No overlap here, corresponding to cells that do not overlap in space, are counted as false positives. The summation of the negative similarity values then gives the total noMissedCells and noFalsePositives.

### Optical Proteomics

The total intracellular protein copy number was determined by quantifying molecules in two distinct populations: discrete, bright foci (e.g., oligomers or clusters) and a diffusely distributed pool. All analyses were performed using custom MATLAB scripts.

The number of molecules aggregated in bright foci was calculated by identifying local maxima in the fluorescence image, summing their integrated intensities, and normalising this sum by the characteristic intensity of a single fluorophore I_single_.

The concentration of the diffuse molecular pool was quantified by fitting the experimental fluorescence data to a physical model of the cell, thereby deconvolving the effects of optical blurring. To prevent the high intensity of foci from biasing this measurement, pixel regions corresponding to detected foci were masked and excluded from the fitting process. A simulated image was generated based on an appropriate geometric model of the cell (e.g., rod or sphere) assuming a uniform fluorophore distribution. This 3D model was projected into 2D and convolved with the microscope’s point-spread function (PSF). The concentration of the diffuse pool was then determined using a linear least-squares method to find the scaling factor that best matched this simulated image to the background-corrected experimental data. The total number of molecules in the pool was the product of this concentration and the geometric volume of the model.

The specific geometric model was chosen to match the cell type. Rod-shaped cells were modelled as a cylinder with hemispherical caps. A single molecular concentration was determined for the entire cell volume. Spherical eukaryotic cells were modelled as two concentric spheres representing the cell boundary and the nucleus. This approach allowed for the simultaneous fitting of two distinct concentrations for the nuclear and cytoplasmic pools. For this compartmental model, the number of molecules counted in foci was also corrected for the microscope’s limited depth-of-field by extrapolating the spot density measured in the focal plane to the full volume of the relevant compartment.

### Omero Integration

MaCADEMIA is compatible with data stored on Omero. Through Open Microscopy Environment functions built into MATLAB, image data can be directly downloaded into MATLAB workspace and MaCADEMIA run.

## Supplementary

**Supplementary figure 1:**
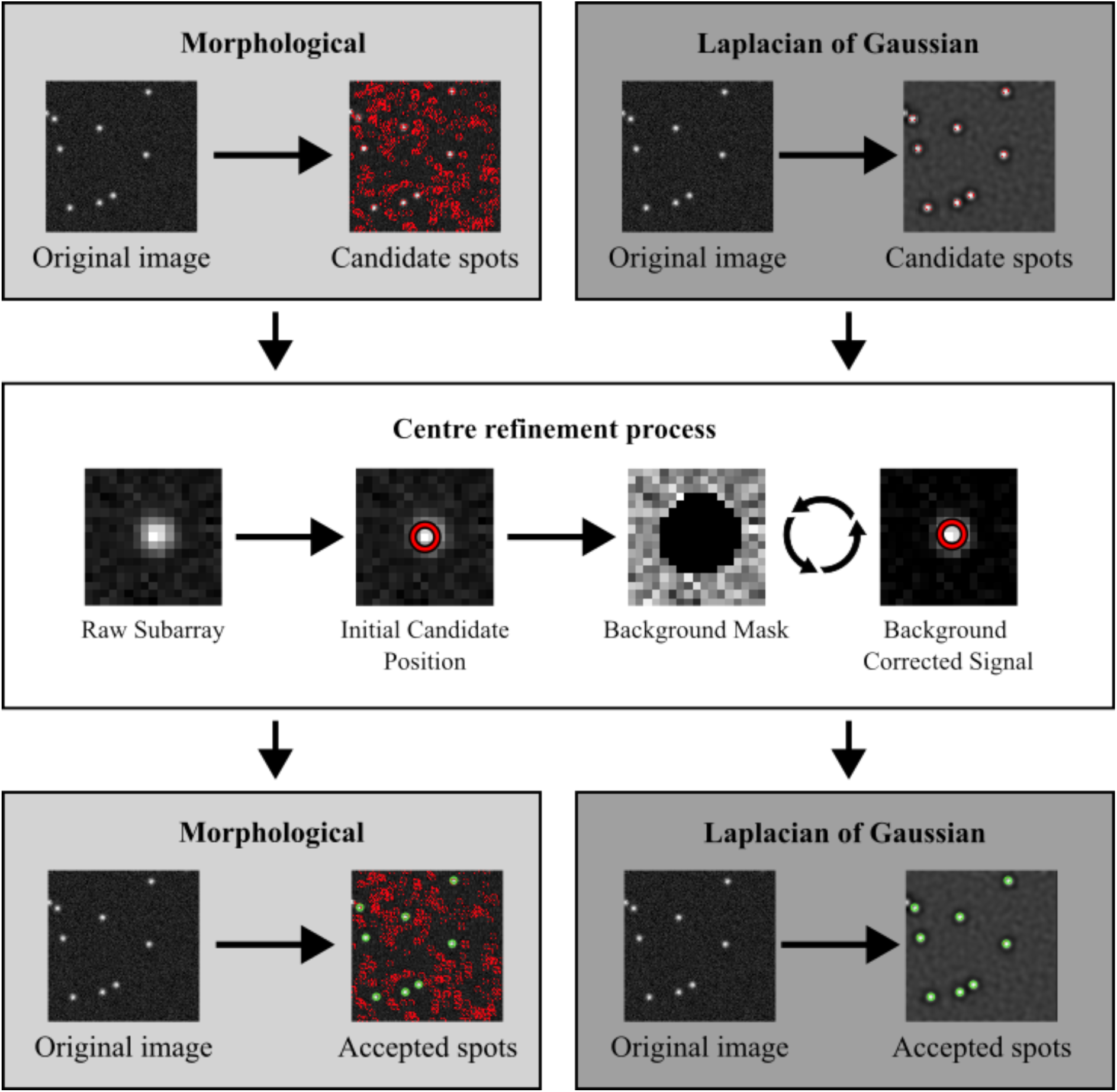
MaCADEMIA single-particle tracking algorithm workffow. A multi-step detection pipeline begins with Candidate Detection, where an adaptive intensity threshold separates signal from background to identify potential spots in the original image. In the Centre Refinement Process, a background mask is generated, and the intensity centroid is calculated to localise each spot with sub-pixel precision. Finally, SNR Postprocessing sums pixel values in the immediate vicinity, subtracts local background, and filters candidates to produce the final set of accepted spots.

**Supplementary figure 2:**
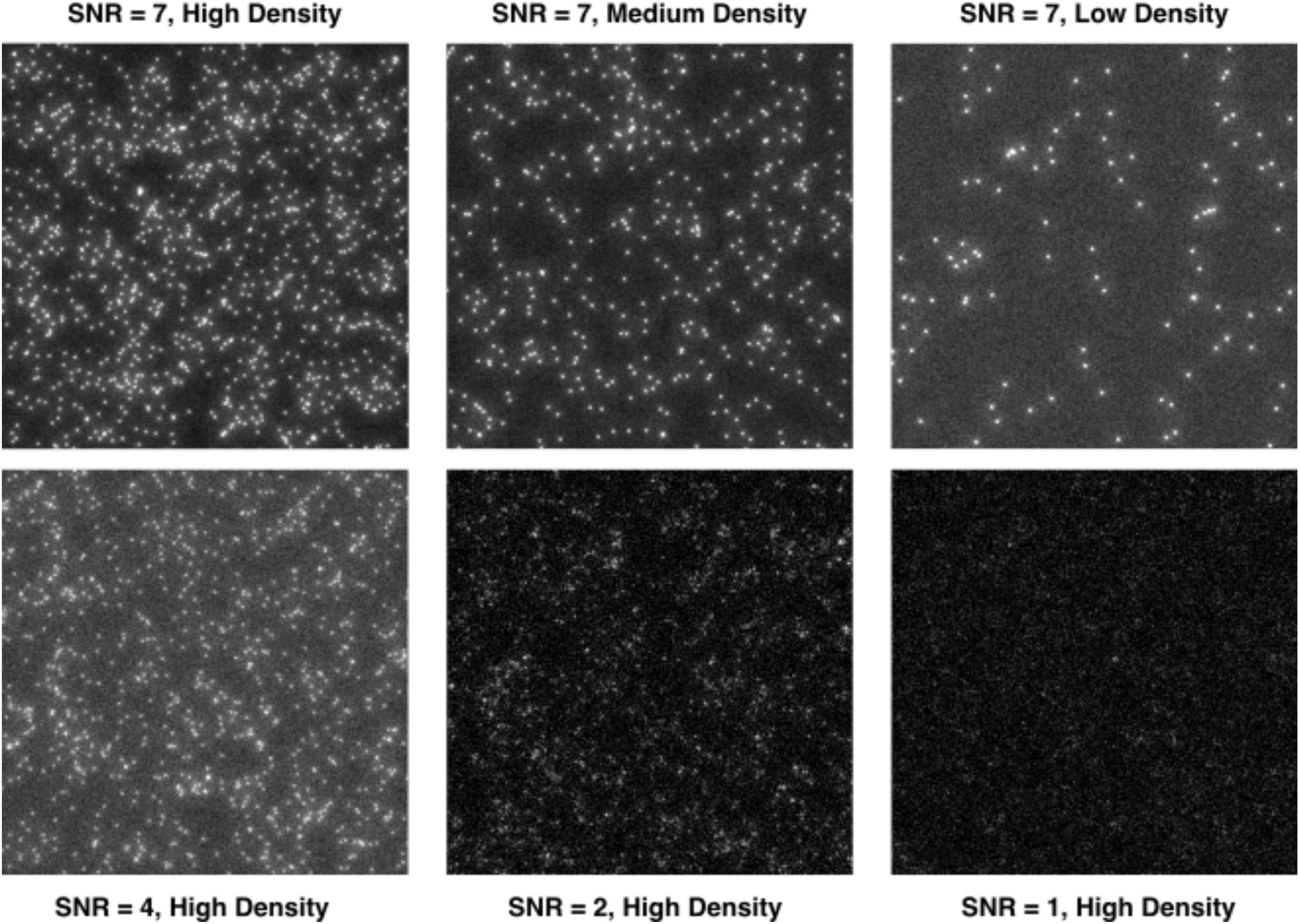
Representative frames from vesicle simulation datasets. Single frames from simulated vesicle datasets demonstrating the range of imaging conditions used for algorithm benchmarking. Top row shows SNR=7 conditions at high, medium, and low particle densities (left to right), illustrating the effect of varying particle concentration on image complexity. Bottom row shows high-density conditions at decreasing SNR values (SNR=4, 2, and 1, left to right), demonstrating progressive degradation of signal quality. At SNR=1, individual particles are barely distinguishable from background noise. Each frame is 512×512 pixels.

**Supplementary figure 3:**
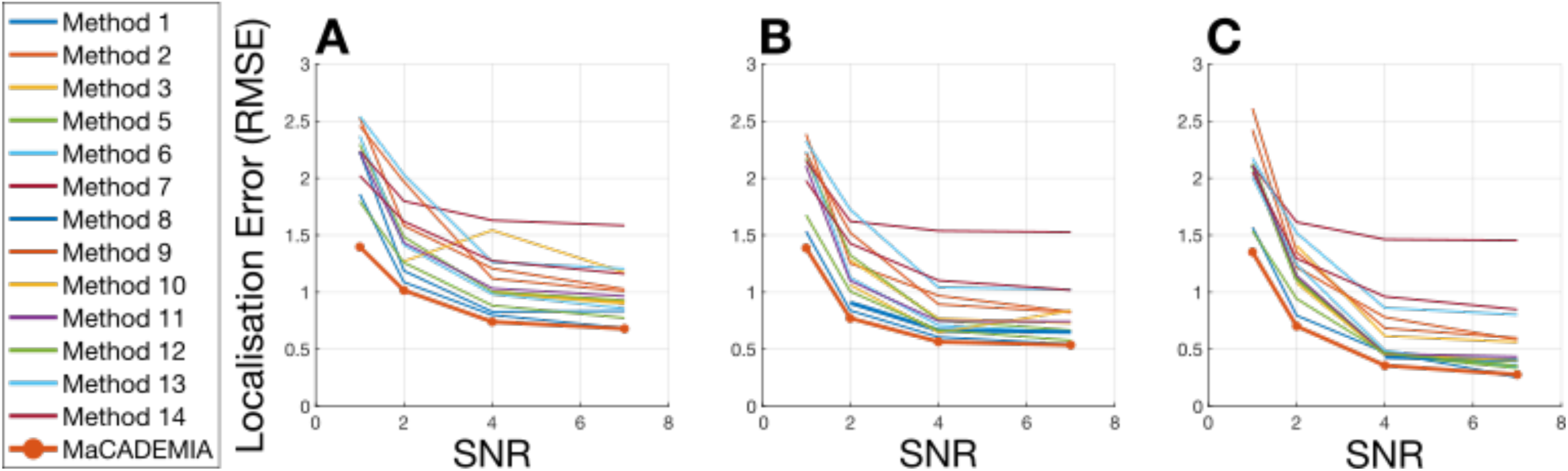
Comparative localisation performance for receptor datasets. Root Mean Square Error (RMSE) of particle localisation plotted against signal-to-noise ratio for receptor simulation datasets at **(A)** high, **(B)** medium, and **(C)** low particle densities. MaCADEMIA (orange line with circles) is compared against 14 peer-reviewed particle tracking algorithms (coloured lines). MaCADEMIA consistently achieves sub-pixel localisation accuracy (RMSE <1 pixel) at SNR ≥4 across all density conditions, placing it among the top-performing methods. At SNR=1, RMSE increases to 1.3-1.5 pixels, comparable to other leading algorithms under these challenging conditions. Performance is most variable at high particle density (A) where proximity effects complicate localisation, but MaCADEMIA maintains competitive accuracy. These results complement Figure 2D-F (vesicle datasets) and demonstrate algorithm robustness across different particle motion models (confined vs. Brownian diffusion).

**Supplementary figure 4:**
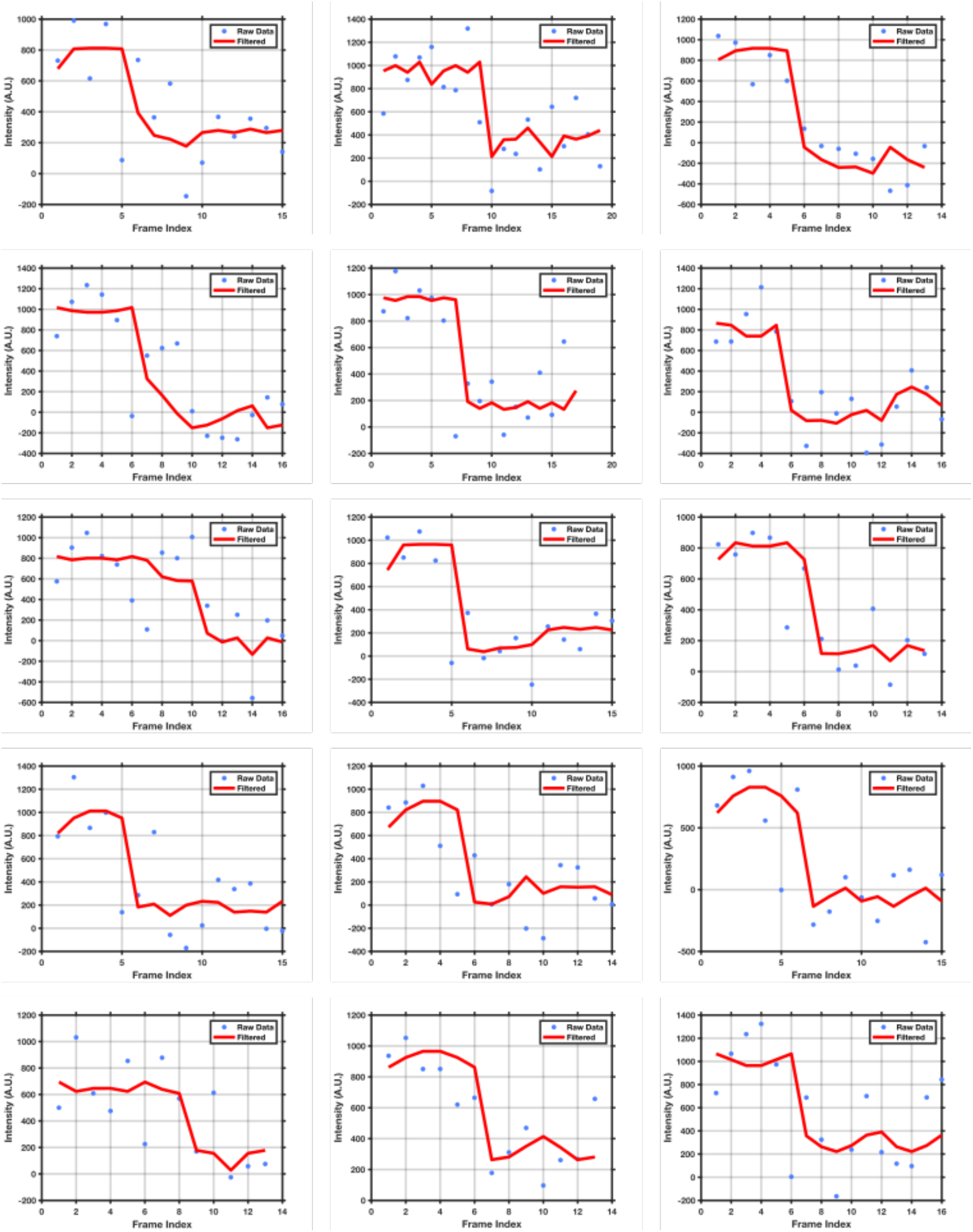
Representative photobleaching trajectories with Chung-Kennedy filtering. Single-molecule intensity trajectories showing raw data (blue points) and Chung-Kennedy filtered traces (red lines, bandwidth = 5, weighting factor = 50). The edge-preserving filter smooths noise while preserving photobleaching steps, enabling stoichiometry determination.

**Supplementary figure 5:**
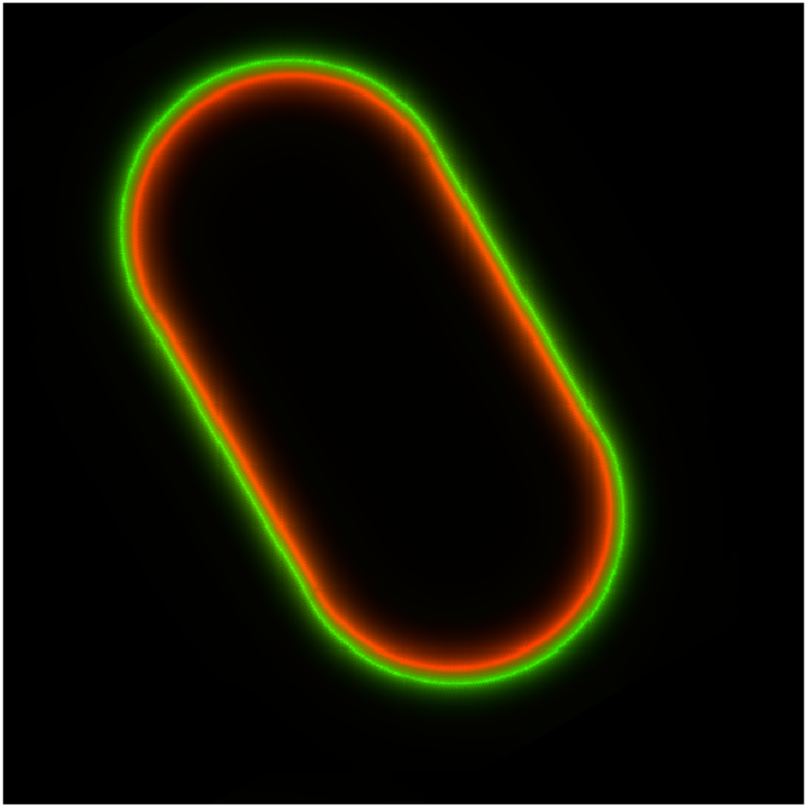
Simulated image of C. elegans membrane and cortex. Membrane in green, simulated as 1 pixel wide and cortex in red simulated as 2 pixels wide using a simulated point spread function approximating a 100x 1.49NA objective lens and real pixel size of 100nm.

